# Multi-omic data integration and analyses for biomarker discovery of spontaneous preterm birth phenotypes

**DOI:** 10.1101/2025.01.31.635906

**Authors:** Juhi K. Gupta, Angharad Care, Laura Goodfellow, Zarko Alfirevic, Ana Alfirevic, Bertram Müller-Myhsok

**Affiliations:** Wolfson Centre for Personalised Medicine, Department of Pharmacology and Therapeutics, Institute of Systems, Molecular and Integrative Biology, University of Liverpool, Liverpool, L69 3GL; Harris-Wellbeing Research Centre, University Department, Liverpool Women’s Hospital, Liverpool, L8 7SS, UK; Max Planck Institute of Psychiatry, 80804, Munich, Germany

**Keywords:** Preterm birth, biomarkers, multi-omics, integrative omics, genomics, transcriptomics, metabolomics

## Abstract

**Background:** Preterm birth, delivery <37 weeks of gestation, is a global health concern affecting 1.3 million infants annually. A higher morbidity burden is associated with infants born <34 weeks of gestation. A more robust biomarker of spontaneous preterm birth needs to be identified. To our knowledge, this study uniquely integrated three ‘ome-wide’ datasets prospectively collected from the same individuals within a single, defined UK cohort. The study aimed to integrate genomic, transcriptomic, and metabolomic data collected from the same cohort of women for biomarker discovery.

**Methods:** Pregnant women with a history of a previous spontaneous preterm birth (sPTB) <34 weeks were recruited in Liverpool in a subsequent pregnancy at 16 and/or 20 weeks of gestation. Pregnancy outcomes were followed up and categorised into the different clinical subgroups of SPTB <34 weeks, PPROM <34 weeks and term delivery >37 weeks (controls). Blood samples were profiled for genomics, transcriptomics and metabolomics. ANOVA analyses were performed at each gestational timepoint. Network enrichment analysis was performed on significant genes.

**Results:** After multi-omic data integration, 43 women at week 16 and 40 women at week 20 of gestation had all three genomics, transcriptomics and metabolomics profiles available. Multiple significant transcripts were detected (p<0.05) from the ANOVA analyses, though these were mostly in non-coding regions.

**Conclusion:** Three different omic data, genomics, transcriptomics and metabolomics, were integrated across the same individuals for biomarker analyses. Molecular signatures were detected that could lead to understanding the pathways involved in the different subgroups of preterm birth: SPTB and PPROM.

## INTRODUCTION

The delivery of an infant before 37 weeks of gestation is defined as preterm birth (PTB) by the World Health Organisation [1]. Preterm birth is the leading cause of mortality in children under the age of 5 and can also severely impact the health of infants that survive [2]. Infants born preterm are at higher risk of suffering from long-term conditions such as motor impairment, neurological conditions and reduced lung capacity amongst other morbidities [3, 4]. One of the main aims of the UK Government’s ‘National Maternity Safety Ambition’ was to reduce PTB rate to 6% by 2025 [5]. However, this target was not met. The Office for National Statistics reported an increase in PTB rate to 7.9% in 2022, showing that progress plateaued [6]. This prompted calls for revised national targets and a renewed focus on evidence-based interventions to reduce preterm birth. Early identification of pregnancies at risk of PTB prior to attending healthcare services with symptoms of labour is a key way we may reduce PTB, allowing for use of preventative interventions.

Many studies define spontaneous PTB as a single timepoint in pregnancy including different clinical presentations of PTB within the same analysis, making it less likely that we will identify a single biomarker. In clinical practice approximately a third of spontaneous preterm births are preceded by preterm prelabour rupture of membranes (PPROM) and do not deliver the baby until days, weeks or even months later, with an increased association with infection [7, 8]. The other two thirds present with more typical signs and symptoms of labour resulting in delivery of the baby. This study will focus on isolated spontaneous preterm births (SPTB) and those preceded by PPROM as two separate groups to identify predictive biomarkers [9].

The term ‘omics’ encompasses multiple types of biological data including genomics, transcriptomics, proteomics, metabolomics and many more, whereby ‘big data’ are generated from high-throughput biotechnologies [10]. Successful integration of multiple omic data sources can enhance our understanding of the relationships between molecules within dynamic biological networks [10, 11, 12]. Analysis of multi-omic data with clinical data could provide insights of cellular functions that are difficult to determine in single omic investigations, especially for complex conditions [11].

However, many challenges arise from ‘big data’ studies including those utilising multi-omic data. The lack of interpretability of the data or results from multivariate analyses and missing data (such as lack of knowledge of non-coding regions of the genome or study participants missing clinic visits) are ongoing challenges; this is further complicated by the availability of bioinformatic tools with the capacity to analyse multi-omic data [13, 14, 15].

Many single omic studies were previously identified in a systematic review by Gupta *et al*., (2022) [16] that aimed to determine biomarkers of PTB, largely proteomics and genomics. Many of these studies identified molecules associated with infection or inflammation pathways, though few of them attempted to combine two different omic datasets. A multi-centre study by Tarca *et al*., (2021) acquired PTB blood biomarker data for transcriptomics and proteomics, determined expression changes in maternal blood samples prior to 33 weeks of gestation [17].

Volkmann et al., (2019) [18] outlined the importance of including clinical data (non-omics) in omic studies to evaluate the predictive value of omic predictors. Lifestyle factors, such as BMI or smoking, have previously been associated with increased risk of spontaneous PTB in a European population [19, 20].

Different omic data integration methods have been described in the literature, of which, the simplest form of combining datasets is the ‘concatenation’ approach (early integration), this method combines different omics data into a single dataset [21]. The ‘concatenation’ method enables integration of many omic data types whilst retaining the original data format and is simpler to perform compared to intermediate or late integration, which require transformation of the data often using mathematical models [21]. This study aimed to apply a multi-omic data integration approach to determine biomarkers of SPTB and PPROM respectively.

## METHODS

### Study participants

Women with a singleton pregnancy and history of spontaneous preterm birth or PPROM <34 weeks were recruited at 16 and 20 weeks of gestation at the Liverpool Women’s hospital (between April 2012 and December 2017). These women are all at high risk for preterm birth based on their obstetric history alone. Whole blood samples were collected for genomic, transcriptomic and metabolomic analyses. Women were followed up after delivery and their pregnancy outcome was collected and classified into case or control groups. Pregnancy outcome information was available for all the women included in this study. Women delivering ≥ 37 weeks of gestation were described as ‘HTERM’ (High-risk term). Alternatively, preterm deliveries were classified into SPTB (spontaneous PTB <34 weeks) or PPROM (PPROM ≤ 34 weeks of gestation). High-risk women delivering between 35-37 weeks were excluded from the analysis. Full details of the inclusion and exclusion criteria for this study cohort are available in previous work (Gupta et al., 2022; Gupta et al., 2021) [22, 23].

### Multi-omic data integration and variables

Genomic SNP data from previous genome-wide association study (GWAS), transcriptomic and metabolomic data were collated from pregnant women recruited in the Liverpool PTB cohort [22, 23]. Post-imputation GWAS data did not yield many genome-wide significant SNPs (p-value < 5×10^-8^), therefore a threshold of p < 1×10^-5^ was applied to capture SNPs approaching the genome-wide significance threshold (876 SNPs in total) [22]. All transcripts (normalised and annotated) were obtained from the transcriptomics expression set (n=138745) [22]. Nuclear Magnetic Resonance (NMR) metabolomic spectral bins (n=145 per timepoint) were scaled prior to integration [23].

The three omic datasets and corresponding clinical phenotypes from the same women were concatenated for each gestational timepoints (16 weeks and 20 weeks) prior to any form or analysis or transformation (‘early integration’ or ‘vertical integration’ approach) in R (Figure 1). The final dataset corresponded to the study participants with all three omic data were available for analyses (Figure 1).

**Figure 1.**
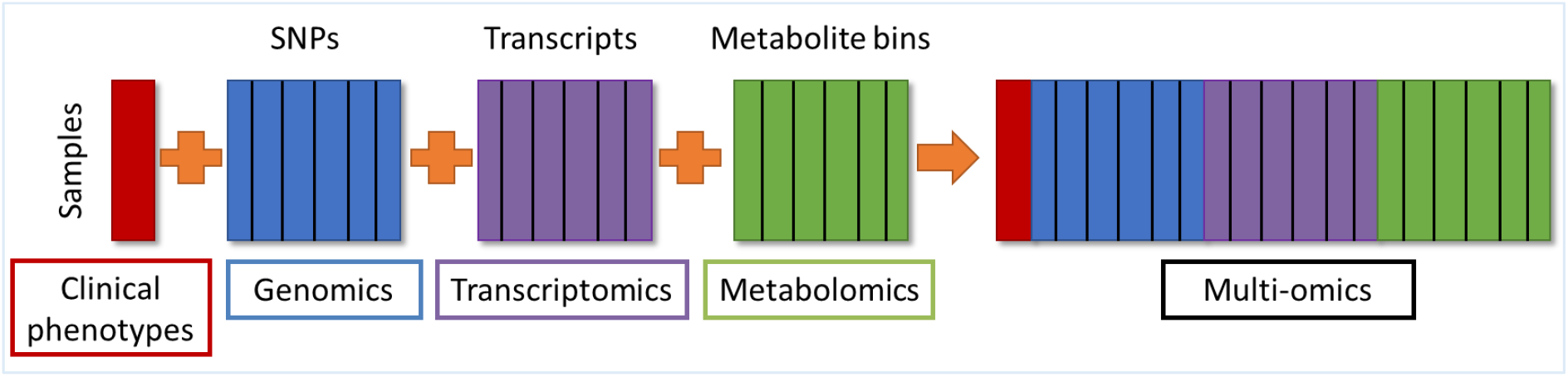
Concatenation (or early integration, or vertical integration) procedure of multiple omic datasets for integrative omic analysis. (SNP = single nucleotide polymorphism)

### Multi-omics significance testing

ANOVA analyses (and Tukey’s HSD post-hoc) performed on the integrated datasets at 16 and 20 weeks of gestation using base R function ‘aov()’. SNPs identified were annotated by applying the Ensembl GRCh37 Biomart data mining tool, and Ensembl gene 101 Human genes (GRCh37.p13). Variables yielding p<0.05 were considered as ‘statistically significant’.

### Network enrichment analysis

To determine the functionality of significant SNPs (with available gene annotations) (p<0.05) resulting from ANOVA analyses, network enrichment analysis was performed. The gene lists from each ANOVA analyses (per gestation) were uploaded to EviNet v1.0 (evidence-based network enrichment analysis) online tool: https://www.evinet.org/ [24]. Network analysis was conducted including input from public databases. Functional gene sets were characterised using KEGG database [25].

## RESULTS

### Multi-omic data integration

Successful integration of genomic, transcriptomic and metabolomic profiles of maternal blood samples was performed for HTERM and both PTB clinical subgroups of SPTB and PPROM. In total, all 3 omic data profiles were available for 43 women at 16 weeks of gestation and 40 women at 20 weeks of gestation. Figure 2 outlines the number of women with multiple omic data available (genomic, transcriptomic and metabolomic) for each clinical phenotype (SPTB, PPROM and HTERM) at week 16 or 20 of gestation.

**Figure 2.**
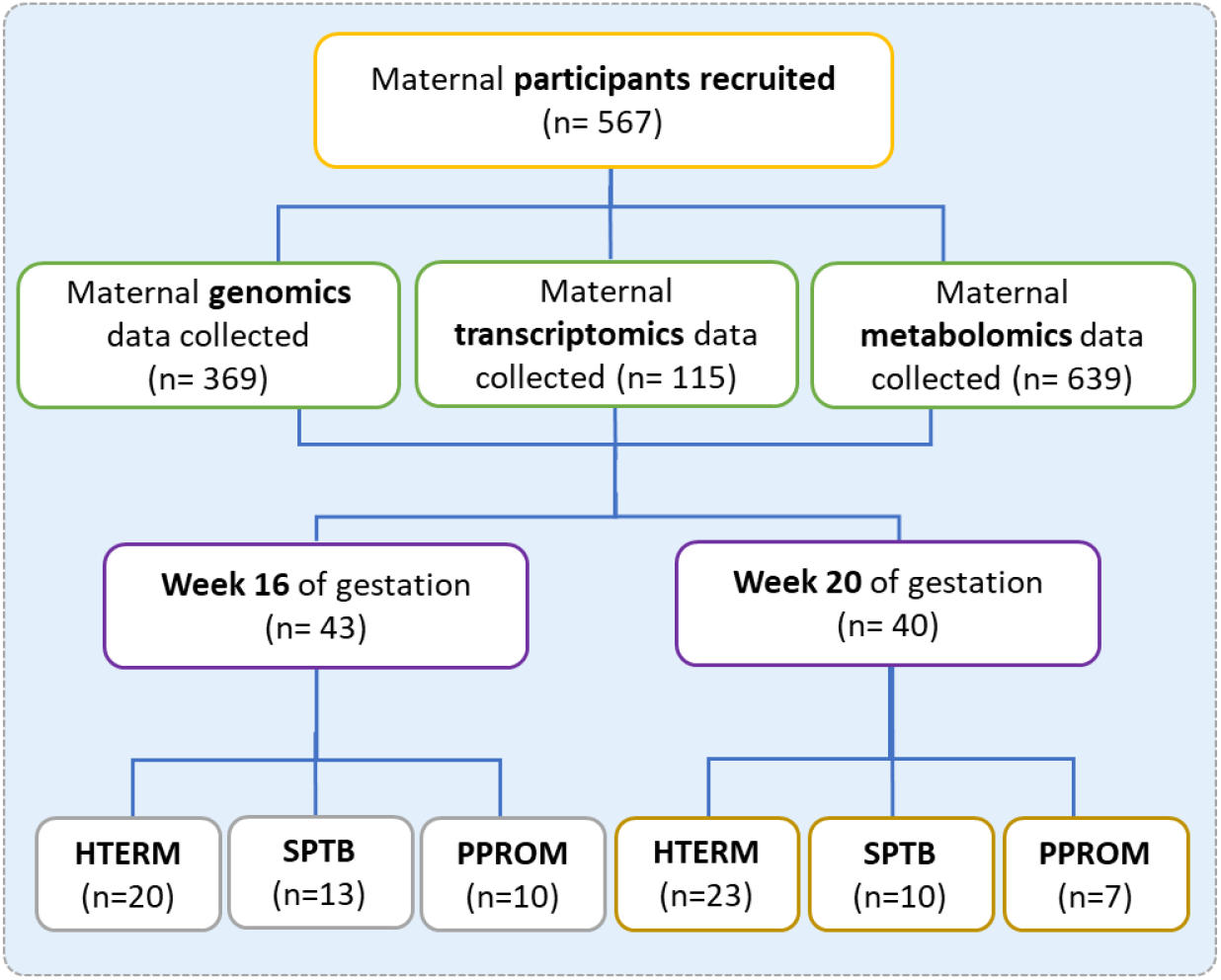
Maternal participants were recruited to the Liverpool PTB cohort (n=567). Data were collected for three different types of omic data (genomics, transcriptomics and metabolomics) for each participant at week 16 (n=43) and week 20 (n=40) of gestation. The number of women with all 3 omic datasets available at both gestational timepoints: HTERM, n=19; PPROM, n=6; SPTB, n=10. A total of 35 women had samples at both week 16 and 20 of gestation and all three omics.

### ANOVA

Statistical analyses of the multi-omic dataset at week 16 of gestation yielded a total of 11,544 omics variables reaching significance at p<0.05, SNPs (n=433), transcripts (n=11,099) and metabolite bins (n=12). Omics variables reaching p<0.0001 (n=37) are summarised in Table 1. Across the phenotype group comparisons, the number of variables that were determined as statistically significant (p<0.05) varied: PPROM-HTERM (n= 5,558), SPTB-HTERM (n= 3,520) and SPTB-PPROM (n= 3,619). Significant SNPs and transcripts were annotated, which were largely mapped to non-coding regions. The annotations for significant SNPs (p<0.05) are summarised in Table S1 (Supplementary File 1)) and for transcripts reaching ANOVA significance of p<0.0001 are outlined in Table S2. Majority of transcripts were in non-coding regions or were of no known consequence (Table S2).

**Table 1.** ANOVA with Tukey’s HSD top variables for week 16 of gestation multi-omic data (ANOVA p<0.0001) (n=37). A total of 11,544 omics variables reached significance p<0.05. Transcripts are represented as transcript cluster IDs.

| Omics Variable | ANOVA | PPROM-HERM | SPTB-HERM | SPTB-PPROM |
| --- | --- | --- | --- | --- |
| TC0500007583.hg.1 | 1.11E-06 | 1.99E-06 | 0.00048 | 0.15011 |
| TC0X00006693.hg.1 | 9.81E-06 | 0.99553 | 2.00E-05 | 0.00019 |
| chr10.12597108_C (rs2296754) | 9.84E-06 | 0.06645 | 0.00101 | 8.77E-06 |
| chr10.12597458_C (rs10906166) | 9.84E-06 | 0.06645 | 0.00101 | 8.77E-06 |
| chr10.12597795_T (rs10906167) | 9.84E-06 | 0.06645 | 0.00101 | 8.77E-06 |
| chr10.12597796_C (rs10906168) | 9.84E-06 | 0.06645 | 0.00101 | 8.77E-06 |
| chr10.12598308_G (rs10906170) | 9.84E-06 | 0.06645 | 0.00101 | 8.77E-06 |
| chr10.12602320_C (rs10906171) | 9.84E-06 | 0.06645 | 0.00101 | 8.77E-06 |
| chr10.12603808_T (rs12764878) | 9.84E-06 | 0.06645 | 0.00101 | 8.77E-06 |
| chr10.12605090_C (rs2648711) | 9.84E-06 | 0.06645 | 0.00101 | 8.77E-06 |
| chr10.12605105_T (rs10906173) | 9.84E-06 | 0.06645 | 0.00101 | 8.77E-06 |
| TC0500007819.hg.1 | 1.12E-05 | 0.00336 | 0.023486 | 6.10E-06 |
| chr4.5024120_A (rs73198536) | 1.13E-05 | 2.06E-05 | 1 | 7.53E-05 |
| chr4.5024391_A (rs55881417) | 1.13E-05 | 2.06E-05 | 1 | 7.53E-05 |
| chr4.5029355_T (rs6828842) | 1.13E-05 | 2.06E-05 | 1 | 7.53E-05 |
| TC0900009423.hg.1 | 1.86E-05 | 0.00615 | 0.020942 | 1.03E-05 |
| TC0100013598.hg.1 | 1.92E-05 | 1.13E-05 | 0.056889 | 0.01568 |
| TC0400012583.hg.1 | 2.17E-05 | 2.30E-05 | 0.007064 | 0.12613 |
| TC0900010785.hg.1 | 2.41E-05 | 1.34E-05 | 0.158414 | 0.00584 |
| TC0900009637.hg.1 | 2.56E-05 | 0.01177 | 2.31E-05 | 0.27598 |
| TC0400009006.hg.1 | 2.91E-05 | 1.64E-05 | 0.190815 | 0.0054 |
| chr2.171332206_T (rs75590771) | 3.35E-05 | 0.64347 | 0.0002 | 0.00013 |
| TC1200007734.hg.1 | 3.42E-05 | 2.53E-05 | 0.599887 | 0.00111 |
| TC0600014102.hg.1 | 3.42E-05 | 0.38348 | 2.13E-05 | 0.00885 |
| TC0900007308.hg.1 | 3.57E-05 | 2.79E-05 | 0.655701 | 0.00096 |
| TC1600010724.hg.1 | 3.69E-05 | 0.82878 | 0.000135 | 0.00022 |
| TC1300007261.hg.1 | 4.04E-05 | 6.85E-05 | 0.999985 | 0.00023 |
| TC1900007159.hg.1 | 4.27E-05 | 3.60E-05 | 0.018169 | 0.08807 |
| TC1000010362.hg.1 | 4.85E-05 | 3.63E-05 | 0.622094 | 0.00138 |
| TC1900007815.hg.1 | 5.10E-05 | 0.33759 | 3.08E-05 | 0.01415 |
| TC0900007167.hg.1 | 5.62E-05 | 0.39814 | 3.52E-05 | 0.01178 |
| TC1600010293.hg.1 | 6.35E-05 | 0.10916 | 3.74E-05 | 0.06592 |
| TC0700012552.hg.1 | 7.55E-05 | 4.76E-05 | 0.413377 | 0.004 |
| TC0500008723.hg.1 | 8.06E-05 | 0.0016 | 0.259518 | 6.50E-05 |
| TC0900006888.hg.1 | 8.24E-05 | 0.00011 | 0.008751 | 0.24967 |
| TC0200010645.hg.1 | 8.63E-05 | 0.00029 | 0.835777 | 0.00019 |
| TC0100015971.hg.1 | 9.06E-05 | 0.11512 | 5.41E-05 | 0.07744 |

Multiple SNPs were identified in intronic regions of genes such as CAMK1D (calcium/calmodulin-dependent protein kinase ID, p=9.84E-06), MYO3B (myosin IIIB, p=3.35E-05) and FOXP1 (forkhead box P1, p=0.0045) (Table S1). Of these significant SNPs, rs1572687 on chromosome 13 was identified as the non-coding microRNA transcript exon variant, MIR5007 (p=0.033) (Table S1).

Twelve metabolite bins also reached significance (p<0.05), but none reached significance at p<0.0001 (Table S3). A total of 6 metabolite bins that have characterised profiles (glucarate (4.14 ppm), glucose (3.90 ppm), glutamate/proline (2.09 ppm), myoinositol (3.58 ppm), proline (2.33 ppm) and 2-hydroxybutyrate (4.02 ppm)) and 6 bins in ‘unknown’ profiles of the spectra (Table S3).

At week 20 of gestation, 7,382 significant omic variables were identified (p<0.05), including: SNPs (n=344), transcripts (n=7008) and metabolite bins (n=30), though only 28 omics variables reached p<0.0001 (n=37) as shown in Table 2. Less omics variables were significant (p<0.05), compared to week 16, between PPROM-HTERM (n= 3,447), SPTB-HTERM (n= 2,080) and SPTB-PPROM (n= 2,214).

**Table 2.** ANOVA with Tukey’s HSD top variables for week 20 of gestation multi-omic data (ANOVA p<0.0001) (n=28). A total of 7,382 omics variables reached significance p<0.05. Transcripts are represented as transcript cluster IDs.

| Omics Variable | ANOVA | PPROM-HTERM | SPTB-HTERM | SPTB-PPROM |
| --- | --- | --- | --- | --- |
| chr4.5024120_A (rs73198536) | 1.02E-09 | 1.19E-09 | 1 | 2.67E-08 |
| chr4.5024391_A (rs55881417) | 1.02E-09 | 1.19E-09 | 1 | 2.67E-08 |
| chr4.5029355_T (rs6828842) | 1.02E-09 | 1.19E-09 | 1 | 2.67E-08 |
| chr1.147119257_A (rs137987097) | 1.09E-06 | 1.16E-06 | 1 | 1.25E-05 |
| TC1000009465.hg.1 | 1.12E-06 | 1.04E-06 | 0.00964 | 0.01232 |
| TC0100012239.hg.1 | 6.48E-06 | 0.00022 | 8.62E-05 | 0.96857 |
| chr10.12597108_C (rs2296754) | 7.14E-06 | 0.09444 | 0.00015 | 1.18E-05 |
| chr10.12597458_C (rs10906166) | 7.14E-06 | 0.09444 | 0.00015 | 1.18E-05 |
| chr10.12597795_T (rs10906167) | 7.14E-06 | 0.09444 | 0.00015 | 1.18E-05 |
| chr10.12597796_C (rs10906168) | 7.14E-06 | 0.09444 | 0.00015 | 1.18E-05 |
| chr10.12598308_G (rs10906170) | 7.14E-06 | 0.09444 | 0.00015 | 1.18E-05 |
| chr10.12602320_C (rs10906171) | 7.14E-06 | 0.09444 | 0.00015 | 1.18E-05 |
| chr10.12603808_T (rs12764878) | 7.14E-06 | 0.09444 | 0.00015 | 1.18E-05 |
| chr10.12605090_C (rs2648711) | 7.14E-06 | 0.09444 | 0.00015 | 1.18E-05 |
| chr10.12605105_T (rs10906173) | 7.14E-06 | 0.09444 | 0.00015 | 1.18E-05 |
| chr3.5925036_C (rs73008350) | 2.37E-05 | 1.71E-05 | 0.86576 | 0.00042 |
| chr8.92813658_C (rs72669709) | 2.50E-05 | 0.51079 | 1.45E-05 | 0.0094 |
| TC0900009097.hg.1 | 3.01E-05 | 0.00022 | 0.27139 | 3.00E-05 |
| TC1200009420.hg.1 | 4.95E-05 | 0.03519 | 4.86E-05 | 0.28911 |
| TC1900007132.hg.1 | 5.59E-05 | 0.00076 | 0.15068 | 4.15E-05 |
| TC0800012165.hg.1 | 5.72E-05 | 0.00012 | 0.76274 | 0.00013 |
| TC0700008043.hg.1 | 6.82E-05 | 0.08881 | 5.06E-05 | 0.15679 |
| TC1400009036.hg.1 | 7.53E-05 | 0.00161 | 0.00055 | 0.99206 |
| TC2000006635.hg.1 | 7.98E-05 | 0.0013 | 0.00073 | 0.96303 |
| TC1000009476.hg.1 | 8.46E-05 | 5.88E-05 | 0.12027 | 0.02602 |
| TC0700006476.hg.1 | 8.60E-05 | 5.08E-05 | 0.29692 | 0.00866 |
| TC1700009262.hg.1 | 8.85E-05 | 5.40E-05 | 0.6197 | 0.00275 |
| TC2200008810.hg.1 | 9.45E-05 | 0.00046 | 0.00297 | 0.61222 |

Gene annotations of significant SNPs at p<0.05 are described in Table S4 (Supplementary file 1) and Table S5 for transcripts reaching significance of p<0.0001 from the ANOVA analysis. Several significant SNPs (p<0.05) were identified in genes: CAMK1D (calcium/calmodulin-dependent protein kinase ID, p=7.14E-06), MYO3B (myosin IIIB, p=0.002324), AKAP6 (A kinase (PRKA) anchor protein 6, p=0.005752) and DNER (delta/notch-like EGF repeat containing, p=0.007503). Again, the non-coding exon transcript, MIR5007, was identified (p=0.046585) (Table S4). Annotation of significant transcripts (p<0.0001) determined that many were either non-coding or there was no known associated consequence (Table S5).

More significant metabolite bins (p<0.05) were identified at week 20 compared to week 16. From a total of 30 metabolites, 13 were located in unknown bins and 17 in characterised bins, including multiple glucose bins, mobile lipids, creatinine (4.06 ppm) and lactate (1.33 ppm) (Table S6).

### Network enrichment analysis

For week 16, a total of 146 genes (from SNP annotations) were uploaded to EviNet from week 16 SNP annotation, mapping 975 subnetworks of genes to pathways Figure S1 (Supplementary File 1). COL5A1 (collagen, type V, alpha 1) (rs11103520, ANOVA p=0.00438) was enriched in the extracellular matrix-receptor interaction KEGG pathway (highest confidence of network enrichment score = 37.9) (Figure Other significant pathways identified, with lower confidences scores, included: vascular smooth muscle contraction (HSA04270) and other inflammation/infection pathways (HSA04750, HSA04657 - IL17 and HSA04668 – TNF signalling).

From week 20 data, 98 genes were uploaded to EviNet from ANOVA SNP annotation, which mapped 782 subnetworks of genes to pathways Figure 4. Week 20 network enrichment analysis also identified extracellular matrix-receptor interaction pathway (KEGG: HSA04512) subnetwork of genes with the highest confidence scores (37.9) Figure S2 (Supplementary File 1). All the shortlisted pathways identified at week 20 were also identified in the week 16 findings.

## DISCUSSION

Unique to this cohort, three different types of ‘ome-wide’ datasets (genomic, transcriptomic and metabolomic) profiled for the same cohort of women were successfully integrated. The number of participants with single omic profiles reduced after integrating all three omic datasets, from original 567 participants down to 43 women at week 16 of gestation and 40 women at week 20 of gestation. Of these, three sets of omics data were available for 35 mothers at both gestational time points. The low number of remaining samples was a limitation for further analyses to distinguish between the pathophysiology of PPROM and SPTB. However, this research has demonstrated that multiple omic data types can be integrated for the same participants recruited via a prospective, nested case-control cohort study.

ANOVA analyses highlighted many significant variables at both week 16 (n=11,544) and 20 of gestation (n=7,382), particularly highlighting differences between PTB cases (SPTB and PPROM) and term births (HTERM) (Table 1; Table 2). The top transcripts identified were non-coding. However, MIR5007 was significant at both timepoints (rs1572687, p<0.05). Further investigation of microRNAs could aid determination of their role in spontaneous PTB. At week 16 of gestation, unknown (4.40 ppm) and glucarate (4.14 ppm), and at week 20, metabolite bins including creatinine (4.06 ppm), lactate (1.33 ppm), 2-hydroxyvalerate/arginine (1.62 ppm), mobile-lipids (1.29 ppm) were significant in previous metabolomics investigations [23].

Network enrichment analysis identified many pathways that could be involved in initiating the onset of early labour. Extracellular matrix-receptor interaction pathway obtained the highest confidence score at both gestation timepoints (Figure 3; Figure 4), in addition to this, the vascular smooth muscle contraction pathway (HSA04270) and other inflammation/infection pathways, involving interleukin-17 (IL17) and tumour necrosis factor (TNF) signalling (HSA04750, HSA04657 and HSA04668). These were identified at two different gestational timepoints, without a validation cohort, implying a key role in early pregnancy. These findings were consistent with smooth muscle contraction versus infection between the different spontaneous PTB phenotypes as described in Capece *et al*., (2014) [26], but in this study we were not able to differentiate potential mechanisms between SPTB and PPROM due to low sample numbers. Furthermore, inflammation markers, such as TNF and IL, have been previously highlighted in PTB literature [26].

**Figure 3.**
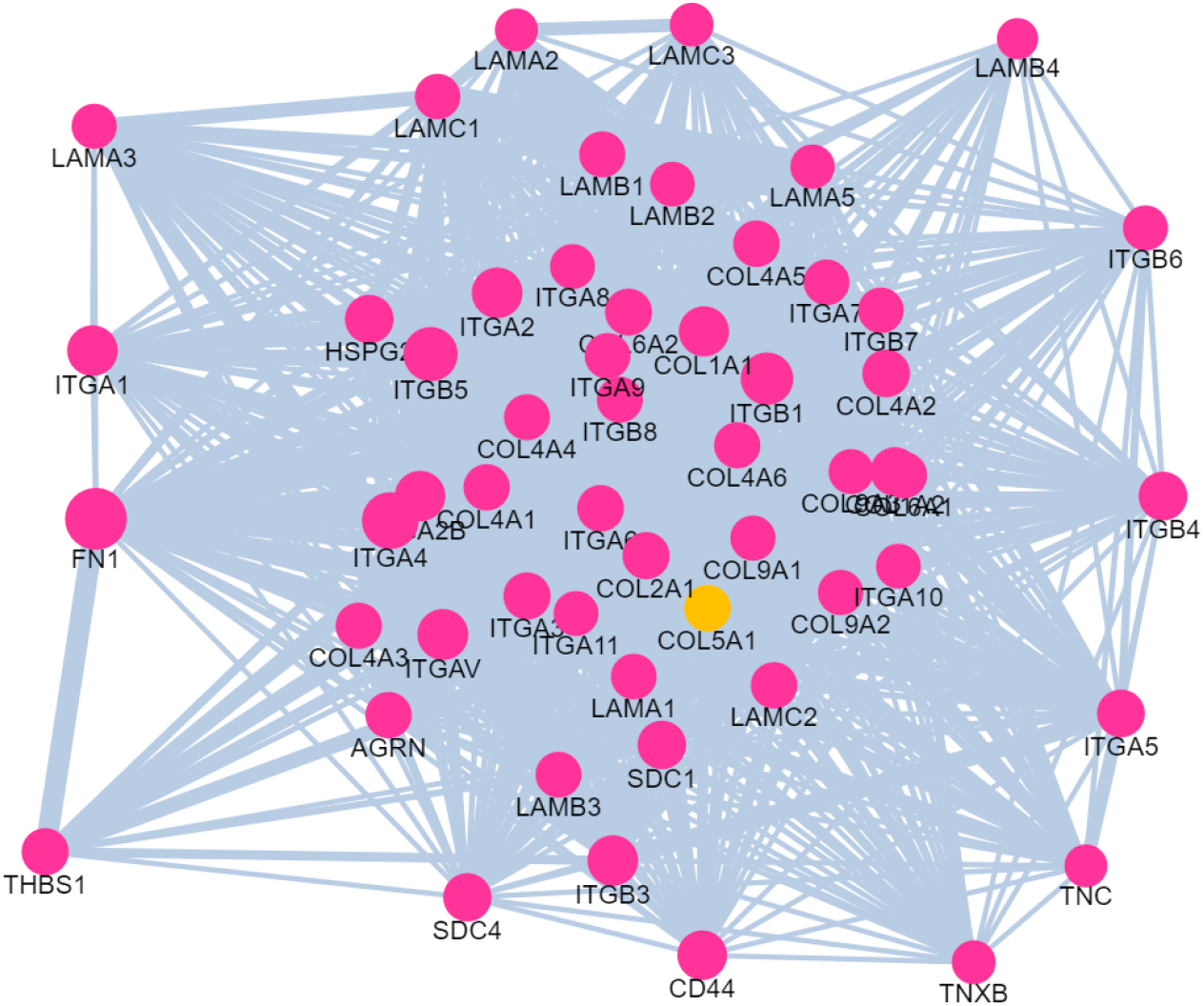
Subnetwork of extracellular matrix-receptor interaction (KEGG: HSA04512) (highest confidence score of 37.9) in association with gene COL5A1 (collagen, type V, alpha 1) identified from ANOVA analysis at week 16 of gestation (rs11103520, p=0.00438). EviNet [24]. Yellow dot = input gene set; pink dot = functional gene sets; edge thickness denotes confidence.

**Figure 4.**
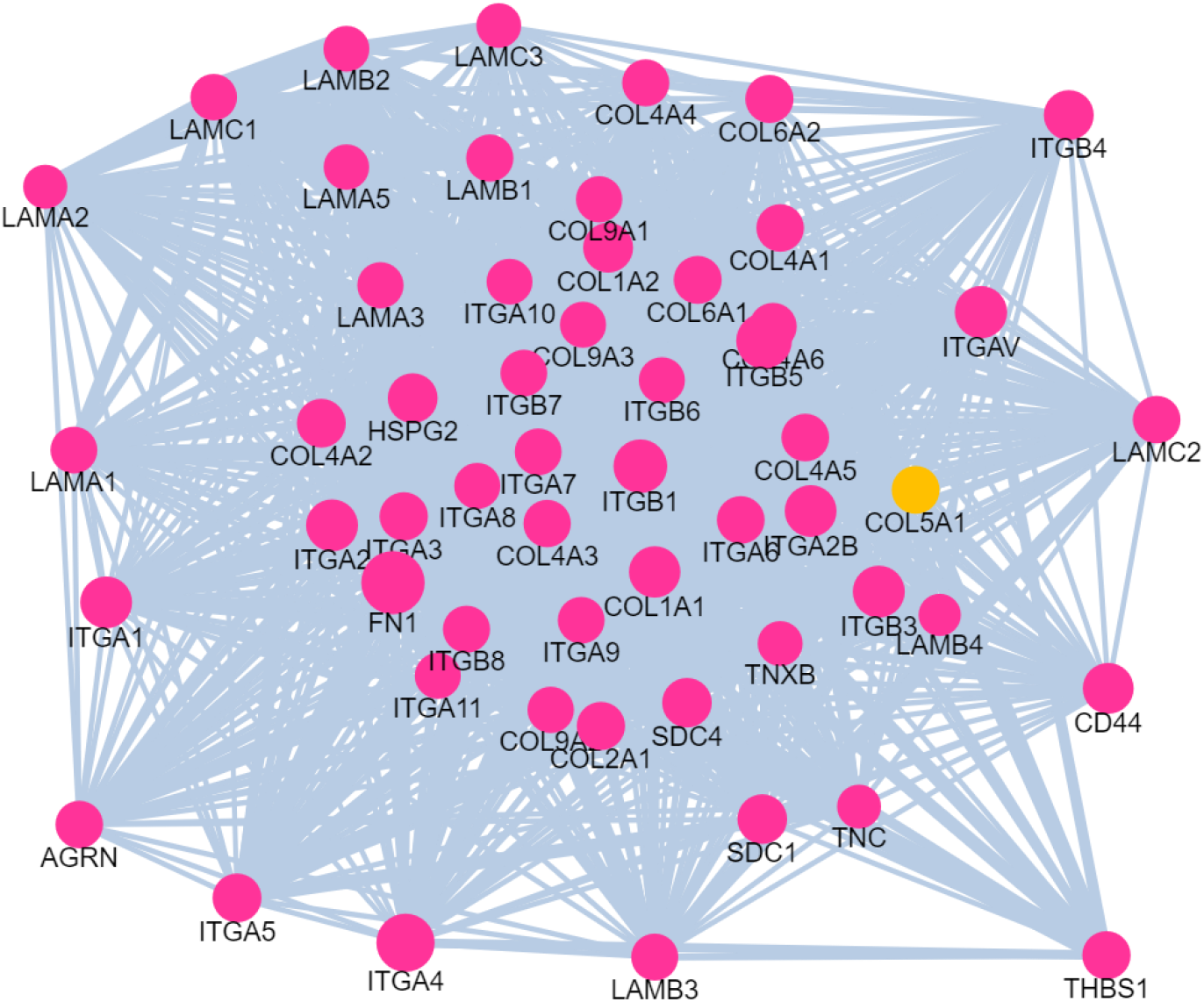
Subnetwork identified in week 20 of gestation network enrichment analysis using EviNet [24]. Extracellular matrix-receptor interaction (KEGG: HSA04512) was highlighted from the analysis. Yellow dot = input gene set; pink dot = functional gene sets; edge thickness denotes confidence.

Jehan *et al*., (2020) [27] also investigated 3 omic data types (transcriptomics, proteomics and metabolomics) from 5 PTB maternal cohorts (defining PTB outcome as < 37 weeks of gestation) from low and middle-income countries [27]. They reported inflammation was associated with PTB as well as metabolism pathways including glutamate, which was also detected in our study and in our previous work [23]. Our study design differed in that we applied multi-omics investigations a prospective, single, well-defined cohort and sampled at two gestation time points, early in pregnancy. We also included genomic data, which enabled us to detect of potential genetic variants that influence the phenotypes of spontaneous PTB.

In our study, all three types of omic datasets were collected from blood samples of the same women that were recruited in Liverpool at two different gestation time points, highlighting this as a unique aspect of this study. Untargeted, ‘ome-wide’ data was acquired and integrated for the two timepoint separately, with 43 participants at week 16 of gestation and 40 at week 20 of gestation. Analyses of omic variables across the different ‘omic layers’ enhanced our understanding of systems biology in spontaneous PTB, which would not be possible with analysis of single omic data alone. Regulation processes occur across multiple omic layers and not in one omic alone, as reviewed in Capece et al., 2014) [26], therefore the ‘multi-omic’ approach applied in this study was appropriate for this purpose. As further work, other types of omic data will be analysed for this cohort, including proteomics. This will aid investigation of the potential pathways that consist of the molecular signatures identified in this study. Furthermore, the results would also need to be validated in other ethnic groups as the women recruited to this study cohort were predominantly from Caucasian ancestry, particularly as the rate of spontaneous PTB is 1.5-fold higher in women of African ancestry [28].

The Liverpool PTB cohort was recruited in a tertiary hospital setting, therefore provisions for specialist care and pregnancy follow-up were available. As the women were recruited in a subsequent pregnancy, obstetric history was also available, in addition to pregnancy outcome and well-defined phenotyping of the outcome. Such facilities are not uniform across the UK or in other countries, therefore the quality or availability of data in other settings may be limited. Environmental factors, such as lifestyle, will also differ across different regions and must be addressed in any analyses.

This prospective study recruited participants with recurrent singletons pregnancies from preterm (≤ 34 weeks of gestation) and healthy term control groups at two gestational timepoints, which is ideal for biomarker discovery in early pregnancy [29]. Blood samples were collected, in an easy and cost-effective manner. This also allows time to detect pathologies and provide timely medical intervention. The biomarker could be developed into a screening test that could be provided for women at high-risk of spontaneous PTB [30, 31]. A screening test would require appropriate validation and identification of who would most benefit from the screening before it could be integrated into clinical practice.

Multi-omics analyses have unveiled potential biomarkers of spontaneous PTB. In addition to understanding the mechanisms of spontaneous PTB, insights into the pathways involved in the initiation of early labour were highlighted. A multi-omics biomarker panel would allow for an effective non-invasive screening tool for women at high risk of spontaneous PTB and therefore aid clinical management of women during pregnancy.

## DECLARATIONS

### Supplementary information

Supplementary file 1

### Authors’ contributions

JG and BMM performed the analyses and JG prepared the manuscript. AA, ZA, BMM and AC designed the study. AC, LG and ZA recruited the participants to the study. All authors reviewed and edited the manuscript.

## Acknowledgements

We would like to thank our funders Wellbeing of Women, UK (Harris-Wellbeing Research Centre). We would also like to acknowledge the study participants; the NMR Centre for Structural Biology and the Centre for Genomic Research (University of Liverpool) for acquiring the metabolomic and genomic data, respectively. We would also like to thank the Oxford Genomics Centre (Wellcome Centre for Human Genetics) for acquiring the genomic SNP data.

## Ethics approval

Research ethics approval was granted by the North West England Research Ethics Committee (REC reference: 11/NW/0720). Informed consent was obtained from all study participants. This study was conducted in compliance with the 1964 Helsinki Declaration and its later amendments or comparable ethical standards.

## Availability of data and material

The genomic and transcriptomic datasets are available in the European Genome-Phenome Archive (EGA) EBI repository (https://ega-archive.org/studies/EGAS00001005076) and the metabolomic data are available in the open-access repository MetaboLights, study ID: MTBLS1990.

