## Supplementary file 1 for "Multi-omic data integration and analyses for biomarker discovery of spontaneous preterm birth phenotypes"

### Multi-omic data integration and analyses for biomarker discovery of spontaneous preterm birth phenotypes

**Juhi K. Gupta**[1,2]*, Angharad Care[2], Laura Goodfellow[2], Zarko Alfirevic[2], Ana Alfirevic[1,2], Bertram Müller-Myhsok[1, 3]

[1] Wolfson Centre for Personalised Medicine, Department of Pharmacology and Therapeutics, Institute of Systems, Molecular and Integrative Biology, University of Liverpool, Liverpool, L69 3GL
[2] Harris-Wellbeing Research Centre, University Department, Liverpool Women's Hospital, Liverpool, L8 7SS, UK
[3]Max Planck Institute of Psychiatry, 80804, Munich, Germany

#### List of supplementary tables and figures

**Table S1.** Gene annotations of single nucleotide polymorphisms at week 16 of gestation (ANOVA, p<0.05). ---------- 2

**Table S2.** Transcript annotations for RNA transcripts determined in week 16 gestation analysis. ----- 7

**Table S3**. Metabolite bins (n=12) identified in week 16 gestation ANOVA with Tukey's HSD analysis (p<0.05). ---------- 8

**Table S4**. Gene annotations of single nucleotide polymorphisms at week 20 of gestation (ANOVA, p<0.05). ---------- 9

**Table S5**. Transcript annotations for RNA transcripts determined in week 20 gestation analysis. ---- 13

**Table S6**. Metabolite bins (n=30) identified in week 20 gestation ANOVA with Tukey's HSD analysis (p<0.05). ---------- 14

**Figure S1**. Week 16 network of significant ANOVA genes generated using EviNet ---------- 15

**Figure S2**. Week 20 network of significant ANOVA genes generated using EviNet ---------- 16

**Table S1. Gene annotations of single nucleotide polymorphisms at week 16 of gestation (ANOVA, p<0.05).** (bp=base pairs)

| Variant | Chromosome | Position start (bp) | Position end (bp) | Variant consequence | HGNC symbol | Gene description | p |
|---|---|---|---|---|---|---|---|
| rs10906166 | 10 | 12597458 | 12597458 | intron variant; non-coding transcript variant | CAMK1D | calcium/calmodulin-dependent protein kinase ID | 0.00000984 |
| rs10906167 | 10 | 12597795 | 12597795 | intron variant; non-coding transcript variant | CAMK1D | calcium/calmodulin-dependent protein kinase ID | 0.00000984 |
| rs10906168 | 10 | 12597796 | 12597796 | intron variant; non-coding transcript variant | CAMK1D | calcium/calmodulin-dependent protein kinase ID | 0.00000984 |
| rs10906170 | 10 | 12598308 | 12598308 | intron variant; non-coding transcript variant | CAMK1D | calcium/calmodulin-dependent protein kinase ID | 0.00000984 |
| rs10906171 | 10 | 12602320 | 12602320 | intron variant; non-coding transcript variant | CAMK1D | calcium/calmodulin-dependent protein kinase ID | 0.00000984 |
| rs10906173 | 10 | 12605105 | 12605105 | intron variant; non-coding transcript variant | CAMK1D | calcium/calmodulin-dependent protein kinase ID | 0.00000984 |
| rs12764878 | 10 | 12603808 | 12603808 | intron variant; non-coding transcript variant | CAMK1D | calcium/calmodulin-dependent protein kinase ID | 0.00000984 |
| rs2296754 | 10 | 12597108 | 12597108 | intron variant; non-coding transcript variant | CAMK1D | calcium/calmodulin-dependent protein kinase ID | 0.00000984 |
| rs2648711 | 10 | 12605090 | 12605090 | intron variant; non-coding transcript variant | CAMK1D | calcium/calmodulin-dependent protein kinase ID | 0.00000984 |
| rs75590771 | 2 | 171332206 | 171332206 | intron variant; non-coding transcript variant; NMD transcript variant | MYO3B | myosin IIIB | 0.0000335 |
| rs9424165 | 10 | 12607024 | 12607024 | intron variant; non-coding transcript variant | CAMK1D | calcium/calmodulin-dependent protein kinase ID | 0.000217 |
| rs137987097 | 1 | 147119257 | 147119257 | intron variant; missense_variant; non-coding transcript exon variant | ACP6 | acid phosphatase 6, lysophosphatidic | 0.000258 |
| rs1409961 | 10 | 12608676 | 12608676 | intron variant; non-coding transcript variant | CAMK1D | calcium/calmodulin-dependent protein kinase ID | 0.000305 |
| rs6602598 | 10 | 12606465 | 12606465 | intron variant; non-coding transcript variant | CAMK1D | calcium/calmodulin-dependent protein kinase ID | 0.000305 |
| rs7896236 | 10 | 12606281 | 12606281 | intron variant; non-coding transcript variant | CAMK1D | calcium/calmodulin-dependent protein kinase ID | 0.000305 |
| rs2037107 | 2 | 11299473 | 11299473 | intron variant; NMD transcript variant; non-coding transcript exon variant | PQLC3 | PQ loop repeat containing 3 | 0.000419 |
| rs62170724 | 2 | 171353456 | 171353456 | intron variant; non-coding transcript variant; NMD transcript variant | MYO3B | myosin IIIB | 0.000728 |
| rs10504272 | 8 | 59900124 | 59900124 | intron variant | TOX | thymocyte selection-associated high mobility group box | 0.001014 |
| rs111937104 | 8 | 59897223 | 59897223 | intron variant | TOX | thymocyte selection-associated high mobility group box | 0.001014 |
| rs61514480 | 8 | 59909111 | 59909111 | intron variant | TOX | thymocyte selection-associated high mobility group box | 0.001014 |
| rs78300448 | 8 | 59906910 | 59906910 | intron variant | TOX | thymocyte selection-associated high mobility group box | 0.001014 |
| rs11159008 | 14 | 25431791 | 25431791 | intron variant; NMD transcript variant | STXBP6 | syntaxin binding protein 6 (amisyn) | 0.002067 |
| rs17257661 | 14 | 25432870 | 25432870 | intron variant; NMD transcript variant | STXBP6 | syntaxin binding protein 6 (amisyn) | 0.002067 |
| rs8181975 | 14 | 25433327 | 25433327 | intron variant; NMD transcript variant | STXBP6 | syntaxin binding protein 6 (amisyn) | 0.002067 |
| rs4668271 | 2 | 171387405 | 171387405 | intron variant; non-coding transcript variant; NMD transcript variant | MYO3B | myosin IIIB | 0.002278 |

| | | | | | | | |
|---|---|---|---|---|---|---|---|
| rs1108432 | 2 | 171368542 | 171368542 | non_coding_transcript_variant; intron variant; NMD transcript variant | MYO3B | myosin IIIB | 0.002611 |
| rs378359 | 6 | 5123453 | 5123453 | intron variant; NMD transcript variant; non-coding transcript exon variant | LYRM4 | LYR motif containing 4 | 0.003112 |
| rs385481 | 6 | 5125574 | 5125574 | intron variant; NMD transcript variant; non-coding transcript exon variant | LYRM4 | LYR motif containing 4 | 0.003112 |
| rs10189774 | 2 | 171375497 | 171375497 | non_coding_transcript_variant; intron variant; NMD transcript variant | MYO3B | myosin IIIB | 0.003112 |
| rs10195667 | 2 | 171376832 | 171376832 | non_coding_transcript_variant; intron variant; NMD transcript variant | MYO3B | myosin IIIB | 0.003112 |
| rs7585843 | 2 | 171385679 | 171385679 | intron variant; non-coding transcript variant; NMD transcript variant | MYO3B | myosin IIIB | 0.003112 |
| rs7595366 | 2 | 171376648 | 171376648 | intron variant; non-coding transcript variant; NMD transcript variant | MYO3B | myosin IIIB | 0.003112 |
| rs141816505 | 22 | 32525653 | 32525653 | intron variant; non-coding transcript variant | AP1B1P1 | adaptor-related protein complex 1, beta 1 subunit pseudogene 1 | 0.003389 |
| rs150619849 | 6 | 152870083 | 152870083 | intron variant; non-coding transcript variant | SYNE1 | spectrin repeat containing, nuclear envelope 1 | 0.003389 |
| rs138952073 | 5 | 26974554 | 26974554 | intron variant; non-coding transcript variant | CDH9 | cadherin 9, type 2 (T1-cadherin) | 0.003389 |
| rs139638763 | 14 | 73821393 | 73821393 | intron variant; non-coding transcript variant | NUMB | numb homolog (Drosophila) | 0.003389 |
| rs190729287 | 4 | 16050206 | 16050206 | intron variant; NMD transcript variant | PROM1 | prominin 1 | 0.003389 |
| rs2329670 | 5 | 41349485 | 41349485 | intron variant | PLCXD3 | phosphatidylinositol-specific phospholipase C, X domain containing 3 | 0.003469 |
| rs78202288 | 2 | 171361842 | 171361842 | intron variant; non-coding transcript variant; NMD transcript variant | MYO3B | myosin IIIB | 0.004063 |
| rs11103520 | 9 | 137664056 | 137664056 | intron variant | COL5A1 | collagen, type V, alpha 1 | 0.00438 |
| rs11041493 | 11 | 7664710 | 7664710 | intron variant; non-coding transcript variant | PPFIBP2 | PTPRF interacting protein, binding protein 2 (liprin beta 2) | 0.004509 |
| rs113228935 | 3 | 155161226 | 155161226 | intron variant | PLCH1 | phospholipase C, eta 1 | 0.004509 |
| rs139867572 | 3 | 71534682 | 71534682 | intron variant; NMD transcript variant | FOXP1 | forkhead box P1 | 0.004509 |
| rs147398711 | 3 | 71526676 | 71526676 | intron variant; NMD transcript variant | FOXP1 | forkhead box P1 | 0.004509 |
| rs17236331 | 5 | 68683069 | 68683069 | intron variant; non-coding transcript variant; NMD transcript variant | RAD17 | RAD17 homolog (S. pombe) | 0.004509 |
| rs35691615 | 16 | 6495078 | 6495078 | intron variant; non-coding transcript variant | RBFOX1 | RNA binding protein, fox-1 homolog (C. elegans) 1 | 0.005727 |
| rs115442065 | 2 | 75357307 | 75357307 | intron variant | TACR1 | tachykinin receptor 1 | 0.006947 |
| rs147946021 | 2 | 75333893 | 75333893 | intron variant | TACR1 | tachykinin receptor 1 | 0.006947 |
| rs75068664 | 2 | 75326498 | 75326498 | intron variant | TACR1 | tachykinin receptor 1 | 0.006947 |
| rs10867845 | 9 | 72252269 | 72252269 | intron variant | APBA1 | amyloid beta (A4) precursor protein-binding, family A, member 1 | 0.007373 |
| rs7774359 | 6 | 5486254 | 5486254 | intron variant | FARS2 | phenylalanyl-tRNA synthetase 2, mitochondrial | 0.008419 |

| | | | | | | | |
|---|---|---|---|---|---|---|---|
| rs137808 | 22 | 44092396 | 44092396 | intron variant; non-coding transcript variant | EFCAB6 | EF-hand calcium binding domain 6 | 0.009232 |
| rs5764188 | 22 | 44052189 | 44052189 | intron variant; non-coding transcript variant | EFCAB6 | EF-hand calcium binding domain 6 | 0.009232 |
| rs5764189 | 22 | 44052763 | 44052763 | intron variant; non-coding transcript variant | EFCAB6 | EF-hand calcium binding domain 6 | 0.009232 |
| rs2355353 | 2 | 171366808 | 171366808 | intron variant; NMD transcript variant; non-coding transcript exon variant | MYO3B | myosin IIIB | 0.011095 |
| rs2103883 | 1 | 61877378 | 61877378 | intron variant | NFIA | nuclear factor I/A | 0.011869 |
| rs9436640 | 1 | 61873677 | 61873677 | intron variant | NFIA | nuclear factor I/A | 0.011869 |
| rs1567481 | 4 | 36030182 | 36030182 | intron variant; non-coding transcript variant | ARAP2 | ArfGAP with RhoGAP domain, ankyrin repeat and PH domain 2 | 0.016117 |
| rs117001634 | 10 | 24567837 | 24567837 | intron variant | KIAA1217 | KIAA1217 | 0.017181 |
| rs1978315 | 16 | 6337331 | 6337331 | intron variant; non-coding transcript variant | RBFOX1 | RNA binding protein, fox-1 homolog (C. elegans) 1 | 0.021035 |
| rs2281599 | 1 | 234432004 | 234432004 | intron variant | SLC35F3 | solute carrier family 35, member F3 | 0.022226 |
| rs147565642 | 10 | 72203622 | 72203622 | intron variant | NODAL | nodal growth differentiation factor | 0.022351 |
| rs1006740 | 2 | 230471160 | 230471160 | intron variant | DNER | delta/notch-like EGF repeat containing | 0.025125 |
| rs10166515 | 2 | 230460953 | 230460953 | intron variant | DNER | delta/notch-like EGF repeat containing | 0.025125 |
| rs10176645 | 2 | 230463982 | 230463982 | intron variant | DNER | delta/notch-like EGF repeat containing | 0.025125 |
| rs10182101 | 2 | 230461604 | 230461604 | intron variant | DNER | delta/notch-like EGF repeat containing | 0.025125 |
| rs10933305 | 2 | 230468480 | 230468480 | intron variant | DNER | delta/notch-like EGF repeat containing | 0.025125 |
| rs11891200 | 2 | 230460267 | 230460267 | intron variant | DNER | delta/notch-like EGF repeat containing | 0.025125 |
| rs11891909 | 2 | 230471427 | 230471427 | intron variant | DNER | delta/notch-like EGF repeat containing | 0.025125 |
| rs12373570 | 2 | 230460640 | 230460640 | intron variant | DNER | delta/notch-like EGF repeat containing | 0.025125 |
| rs12373571 | 2 | 230460818 | 230460818 | intron variant | DNER | delta/notch-like EGF repeat containing | 0.025125 |
| rs12998709 | 2 | 230460466 | 230460466 | intron variant | DNER | delta/notch-like EGF repeat containing | 0.025125 |
| rs13021776 | 2 | 230460178 | 230460178 | intron variant | DNER | delta/notch-like EGF repeat containing | 0.025125 |
| rs1465346 | 2 | 230470702 | 230470702 | intron variant | DNER | delta/notch-like EGF repeat containing | 0.025125 |
| rs1465347 | 2 | 230470738 | 230470738 | intron variant | DNER | delta/notch-like EGF repeat containing | 0.025125 |
| rs1861616 | 2 | 230462275 | 230462275 | intron variant | DNER | delta/notch-like EGF repeat containing | 0.025125 |
| rs2052300 | 2 | 230461647 | 230461647 | intron variant | DNER | delta/notch-like EGF repeat containing | 0.025125 |
| rs2052303 | 2 | 230461850 | 230461850 | intron variant | DNER | delta/notch-like EGF repeat containing | 0.025125 |
| rs2052304 | 2 | 230462056 | 230462056 | intron variant | DNER | delta/notch-like EGF repeat containing | 0.025125 |
| rs2052306 | 2 | 230464359 | 230464359 | intron variant | DNER | delta/notch-like EGF repeat containing | 0.025125 |
| rs2075256 | 2 | 230462384 | 230462384 | intron variant | DNER | delta/notch-like EGF repeat containing | 0.025125 |
| rs2075257 | 2 | 230462394 | 230462394 | intron variant | DNER | delta/notch-like EGF repeat containing | 0.025125 |
| rs2396694 | 2 | 230462929 | 230462929 | intron variant | DNER | delta/notch-like EGF repeat containing | 0.025125 |
| rs2396695 | 2 | 230462932 | 230462932 | intron variant | DNER | delta/notch-like EGF repeat containing | 0.025125 |
| rs6751880 | 2 | 230463620 | 230463620 | intron variant | DNER | delta/notch-like EGF repeat containing | 0.025125 |
| rs759534 | 2 | 230462463 | 230462463 | intron variant | DNER | delta/notch-like EGF repeat containing | 0.025125 |
| rs759535 | 2 | 230463103 | 230463103 | intron variant | DNER | delta/notch-like EGF repeat containing | 0.025125 |
| rs759536 | 2 | 230463185 | 230463185 | intron variant | DNER | delta/notch-like EGF repeat containing | 0.025125 |

| | | | | | | | |
|---|---|---|---|---|---|---|---|
| rs759537 | 2 | 230463271 | 230463271 | intron variant | DNER | delta/notch-like EGF repeat containing | 0.025125 |
| rs759538 | 2 | 230463394 | 230463394 | intron variant | DNER | delta/notch-like EGF repeat containing | 0.025125 |
| rs759539 | 2 | 230465606 | 230465606 | intron variant | DNER | delta/notch-like EGF repeat containing | 0.025125 |
| rs888176 | 2 | 230461255 | 230461255 | intron variant | DNER | delta/notch-like EGF repeat containing | 0.025125 |
| rs888177 | 2 | 230461323 | 230461323 | intron variant | DNER | delta/notch-like EGF repeat containing | 0.025125 |
| rs2742658 | 1 | 3070998 | 3070998 | intron variant; non-coding transcript variant | PRDM16 | PR domain containing 16 | 0.025993 |
| rs77207912 | 15 | 45789759 | 45789759 | intron variant | SLC30A4 | solute carrier family 30 (zinc transporter), member 4 | 0.026053 |
| rs146115003 | 14 | 93276818 | 93276818 | intron variant; non-coding transcript exon variant | GOLGA5 | golgin A5 | 0.026053 |
| rs141575794 | 17 | 62507740 | 62588982 | intron variant; NMD transcript variant | CEP95 | centrosomal protein 95kDa | 0.029888 |
| rs150957438 | 13 | 72281791 | 72281791 | intron variant | DACH1 | dachshund homolog 1 (Drosophila) | 0.029888 |
| rs188293839 | 7 | 121021414 | 121021414 | intron variant | FAM3C | family with sequence similarity 3, member C | 0.029888 |
| rs191854224 | 13 | 72327804 | 72327804 | intron variant | DACH1 | dachshund homolog 1 (Drosophila) | 0.029888 |
| rs2160419 | 13 | 72314346 | 72314346 | intron variant | DACH1 | dachshund homolog 1 (Drosophila) | 0.029888 |
| rs12188035 | 5 | 76638444 | 76638444 | intron variant | PDE8B | phosphodiesterase 8B | 0.029888 |
| rs34344194 | 12 | 15925995 | 15925995 | intron variant; NMD transcript variant | EPS8 | epidermal growth factor receptor pathway substrate 8 | 0.029888 |
| rs61908138 | 12 | 15946956 | 15946956 | intron variant | EPS8 | epidermal growth factor receptor pathway substrate 8 | 0.029888 |
| rs61908155 | 12 | 16010104 | 16010104 | intron variant | EPS8 | epidermal growth factor receptor pathway substrate 8 | 0.029888 |
| rs2092868 | 1 | 61877393 | 61877393 | intron variant | NFIA | nuclear factor I/A | 0.031006 |
| rs150386235 | 6 | 152839337 | 152839337 | intron variant; non-coding transcript variant | SYNE1 | spectrin repeat containing, nuclear envelope 1 | 0.031108 |
| rs113624632 | 10 | 72137038 | 72137038 | intron variant | LRRC20 | leucine rich repeat containing 20 | 0.031868 |
| rs11594504 | 10 | 72173518 | 72173518 | intron variant | EIF4EBP2 | eukaryotic translation initiation factor 4E binding protein 2 | 0.031868 |
| rs11595603 | 10 | 72140069 | 72140069 | intron variant | LRRC20 | leucine rich repeat containing 20 | 0.031868 |
| rs11597457 | 10 | 72177911 | 72177911 | intron variant | EIF4EBP2 | eukaryotic translation initiation factor 4E binding protein 2 | 0.031868 |
| rs16927606 | 10 | 72185254 | 72185254 | 3_prime_UTR_variant | EIF4EBP2 | eukaryotic translation initiation factor 4E binding protein 2 | 0.031868 |
| rs3829188 | 10 | 72185440 | 72185440 | 3_prime_UTR_variant | EIF4EBP2 | eukaryotic translation initiation factor 4E binding protein 2 | 0.031868 |
| rs77125060 | 10 | 72203094 | 72203094 | intron variant | NODAL | nodal growth differentiation factor | 0.031868 |
| rs78934060 | 10 | 72196589 | 72196589 | intron variant | NODAL | nodal growth differentiation factor | 0.031868 |
| rs79198291 | 10 | 72171149 | 72171149 | intron variant | EIF4EBP2 | eukaryotic translation initiation factor 4E binding protein 2 | 0.031868 |
| rs79436270 | 10 | 72138823 | 72138823 | intron variant | LRRC20 | leucine rich repeat containing 20 | 0.031868 |

| | | | | | | | |
|---|---|---|---|---|---|---|---|
| rs184101006 | 12 | 132234284 | 132234284 | intron variant; non-coding transcript variant; NMD transcript variant | SFSWAP | splicing factor, suppressor of white-apricot homolog (Drosophila) | 0.031868 |
| rs189998603 | 12 | 132268394 | 132268394 | intron variant; NMD transcript variant; non-coding transcript exon variant | SFSWAP | splicing factor, suppressor of white-apricot homolog (Drosophila) | 0.031868 |
| rs72824604 | 17 | 11182840 | 11182840 | intron variant | SHISA6 | shisa family member 6 | 0.032772 |
| rs1572687 | 13 | 55748673 | 55748673 | non-coding transcript exon variant | MIR5007 | microRNA 5007 | 0.032911 |
| rs2742659 | 1 | 3070829 | 3070829 | intron variant; non-coding transcript variant | PRDM16 | PR domain containing 16 | 0.034545 |
| rs62183592 | 2 | 190429752 | 190429752 | intron variant | SLC40A1 | solute carrier family 40 (iron-regulated transporter), member 1 | 0.035515 |
| rs6587924 | 1 | 61895257 | 61895257 | intron variant | NFIA | nuclear factor I/A | 0.036255 |
| rs6587925 | 1 | 61895589 | 61895589 | intron variant | NFIA | nuclear factor I/A | 0.036255 |
| rs1362318 | 16 | 6336762 | 6336762 | intron variant; non-coding transcript variant | RBFOX1 | RNA binding protein, fox-1 homolog (C. elegans) 1 | 0.041379 |
| rs1420036 | 16 | 6337686 | 6337686 | intron variant; non-coding transcript variant | RBFOX1 | RNA binding protein, fox-1 homolog (C. elegans) 1 | 0.041379 |
| rs17220445 | 16 | 6338473 | 6338473 | intron variant; non-coding transcript variant | RBFOX1 | RNA binding protein, fox-1 homolog (C. elegans) 1 | 0.041379 |
| rs17220529 | 16 | 6338548 | 6338548 | intron variant; non-coding transcript variant | RBFOX1 | RNA binding protein, fox-1 homolog (C. elegans) 1 | 0.041379 |
| rs17221054 | 16 | 6342445 | 6342445 | intron variant; non-coding transcript variant | RBFOX1 | RNA binding protein, fox-1 homolog (C. elegans) 1 | 0.041379 |
| rs17819872 | 16 | 6339757 | 6339757 | intron variant; non-coding transcript variant | RBFOX1 | RNA binding protein, fox-1 homolog (C. elegans) 1 | 0.041379 |
| rs17819914 | 16 | 6339784 | 6339784 | intron variant; non-coding transcript variant | RBFOX1 | RNA binding protein, fox-1 homolog (C. elegans) 1 | 0.041379 |
| rs1978314 | 16 | 6337404 | 6337404 | intron variant; non-coding transcript variant | RBFOX1 | RNA binding protein, fox-1 homolog (C. elegans) 1 | 0.041379 |
| rs1978317 | 16 | 6337230 | 6337230 | intron variant; non-coding transcript variant | RBFOX1 | RNA binding protein, fox-1 homolog (C. elegans) 1 | 0.041379 |
| rs4255786 | 16 | 6337914 | 6337914 | intron variant; non-coding transcript variant | RBFOX1 | RNA binding protein, fox-1 homolog (C. elegans) 1 | 0.041379 |
| rs4255787 | 16 | 6338221 | 6338221 | intron variant; non-coding transcript variant | RBFOX1 | RNA binding protein, fox-1 homolog (C. elegans) 1 | 0.041379 |
| rs4255788 | 16 | 6338235 | 6338235 | intron variant; non-coding transcript variant | RBFOX1 | RNA binding protein, fox-1 homolog (C. elegans) 1 | 0.041379 |
| rs4274443 | 16 | 6337919 | 6337919 | intron variant; non-coding transcript variant | RBFOX1 | RNA binding protein, fox-1 homolog (C. elegans) 1 | 0.041379 |
| rs55680138 | 16 | 6340264 | 6340264 | intron variant; non-coding transcript variant | RBFOX1 | RNA binding protein, fox-1 homolog (C. elegans) 1 | 0.041379 |
| rs56193200 | 16 | 6340315 | 6340315 | intron variant; non-coding transcript variant | RBFOX1 | RNA binding protein, fox-1 homolog (C. elegans) 1 | 0.041379 |
| rs67861918 | 16 | 6338425 | 6338425 | intron variant; non-coding transcript variant | RBFOX1 | RNA binding protein, fox-1 homolog (C. elegans) 1 | 0.041379 |
| rs716509 | 16 | 6336737 | 6336737 | intron variant; non-coding transcript variant | RBFOX1 | RNA binding protein, fox-1 homolog (C. elegans) 1 | 0.041379 |
| rs7199065 | 16 | 6343493 | 6343493 | intron variant; non-coding transcript variant | RBFOX1 | RNA binding protein, fox-1 homolog (C. elegans) 1 | 0.041379 |
| rs7202627 | 16 | 6343694 | 6343694 | intron variant; non-coding transcript variant | RBFOX1 | RNA binding protein, fox-1 homolog (C. elegans) 1 | 0.041379 |
| rs5016273 | 1 | 61881512 | 61881512 | intron variant | NFIA | nuclear factor I/A | 0.045954 |
| rs1991611 | 10 | 77380716 | 77380716 | intron variant; non-coding transcript variant | C10orf11 | chromosome 10 open reading frame 11 | 0.048352 |

**Table S2. Transcript annotations for RNA transcripts determined in week 16 gestation analysis.** (bp=base pairs)

| Transcript cluster ID | Chromosome | Start position (bp) | End position (bp) | Gene | Gene name | Locus type | p |
|---|---|---|---|---|---|---|---|
| TC0500007583.hg.1 | chr5 | 63919841 | 63941223 | | | NonCoding | 1.11E-06 |
| TC0X00006693.hg.1 | chrX | 16581281 | 16584295 | | | NonCoding | 9.81E-06 |
| TC0500007819.hg.1 | chr5 | 76078666 | 76079763 | CTC-235G5.2 | | Pseudogene | 1.12E-05 |
| TC0900009423.hg.1 | chr9 | 3770287 | 3814472 | | | NonCoding | 1.86E-05 |
| TC0100013598.hg.1 | chr1 | 32101130 | 32102707 | | | NonCoding | 1.92E-05 |
| TC0400012583.hg.1 | chr4 | 183639635 | 183659225 | RWDD4 | RWD domain containing 4 | Multiple_Complex | 2.17E-05 |
| TC0900010785.hg.1 | chr9 | 92145869 | 92146323 | | | NonCoding | 2.41E-05 |
| TC0900009637.hg.1 | chr9 | 18717974 | 18718526 | RAP1BP1 | RAP1B | Multiple_Complex | 2.56E-05 |
| TC0400009006.hg.1 | chr4 | 151730395 | 151731057 | sterbeebo | Transcript Identified by AceView | Coding | 2.91E-05 |
| TC1200007734.hg.1 | chr12 | 53965965 | 53966061 | HOTAIR_3 | HOTAIR conserved region 3 | NonCoding | 3.42E-05 |
| TC0600014102.hg.1 | chr6 | 31463180 | 31477506 | HCP5 | HLA complex P5 (non-protein coding) | Multiple_Complex | 3.42E-05 |
| TC0900007308.hg.1 | chr9 | 42974855 | 42978320 | GXYLT1P5 | glucoside xylosyltransferase 1 pseudogene 5 | Multiple_Complex | 3.57E-05 |
| TC1600010724.hg.1 | chr16 | 69443240 | 69444290 | RP11-343C2.10 | | Multiple_Complex | 3.69E-05 |
| TC1300007261.hg.1 | chr13 | 53345211 | 53410880 | RP11-384G23.1 | novel transcript | NonCoding | 4.04E-05 |
| TC1900007159.hg.1 | chr19 | 14119106 | 14119537 | CTB-55O6.10 | | NonCoding | 4.27E-05 |
| TC1000010362.hg.1 | chr10 | 37775206 | 37858106 | ZNF248 | zinc finger protein 248 | Multiple_Complex | 4.85E-05 |
| TC1900007815.hg.1 | chr19 | 34594396 | 34676192 | | | NonCoding | 0.000051 |
| TC0900007167.hg.1 | chr9 | 37800502 | 37867668 | DCAF10 | DDB1 and CUL4 associated factor 10 | Multiple_Complex | 5.62E-05 |
| TC1600010293.hg.1 | chr16 | 51180767 | 51181457 | swoyry | Transcript Identified by AceView | Coding | 6.35E-05 |
| TC0700012552.hg.1 | chr7 | 128928191 | 128928571 | kawflawbu | Transcript Identified by AceView | Coding | 7.55E-05 |
| TC0500008723.hg.1 | chr5 | 135449574 | 135458868 | CTB-138E5.1 | | NonCoding | 8.06E-05 |
| TC0900006888.hg.1 | chr9 | 27210327 | 27210626 | RP11-179D22.1 | | Multiple_Complex | 8.24E-05 |
| TC0200010645.hg.1 | chr2 | 210028417 | 210029156 | RP11-260M2.1 | | NonCoding | 8.63E-05 |
| TC0100015971.hg.1 | chr1 | 155749658 | 155859400 | GON4L | gon-4-like (C. elegans) | Multiple_Complex | 9.06E-05 |

**Table S3. Metabolite bins (n=12) identified in week 16 gestation ANOVA with Tukey's HSD analysis (p<0.05).**

Breakdown of metabolite bins with p<0.05 for each phenotypic group comparison: PPROM-HTERM (n=0); SPTB-HTERM (n=11); SPTB-PPROM (n=0).

| Omics variable | ANOVA | PPROM-HTERM | SPTB-HTERM | SPTB-PPROM |
|---|---|---|---|---|
| unknown (3.81ppm) | 0.012876533 | 0.655334138 | 0.00950442 | 0.175847396 |
| unknown (2.58ppm) | 0.017200156 | 0.367813029 | 0.013049842 | 0.419627899 |
| myoinositol (3.58ppm) | 0.020733106 | 0.910628932 | 0.01854587 | 0.117544258 |
| proline (2.33ppm) | 0.030645591 | 0.912135841 | 0.027297253 | 0.150921275 |
| unknown (3.88ppm) | 0.032990432 | 0.298720312 | 0.029109525 | 0.661968321 |
| glucose (3.90ppm) | 0.036987348 | 0.747639208 | 0.029087208 | 0.263332719 |
| unknown (3.80ppm) | 0.038244157 | 0.934990363 | 0.035080022 | 0.160829673 |
| glutamate/proline (2.09ppm) | 0.043935513 | 0.929222604 | 0.039863552 | 0.179744361 |
| unknown (1.41ppm) | 0.045521399 | 0.748539725 | 0.03603946 | 0.29651368 |
| glucarate (4.14ppm) | 0.046713479 | 0.612566906 | 0.036238575 | 0.400683746 |
| unknown (4.40ppm) | 0.047140854 | 0.706874596 | 0.036925947 | 0.331163264 |
| 2-hydroxybutyrate (4.02ppm) | 0.049916234 | 0.739426175 | 0.125945723 | 0.056932817 |

**Table S4. Gene annotations of single nucleotide polymorphisms at week 20 of gestation (ANOVA, p<0.05).** (bp=base pairs)

| Variant name | Chromosome | Position start (bp) | Position end (bp) | Variant consequence | HGNC symbol | Gene description | p |
|---|---|---|---|---|---|---|---|
| rs137987097 | 1 | 147119257 | 147119257 | intron variant; missense_variant; non-coding transcript exon variant | ACP6 | acid phosphatase 6, lysophosphatidic | 1.09E-06 |
| rs10906166 | 10 | 12597458 | 12597458 | intron variant; non-coding transcript variant | CAMK1D | calcium/calmodulin-dependent protein kinase ID | 7.14E-06 |
| rs10906167 | 10 | 12597795 | 12597795 | intron variant; non-coding transcript variant | CAMK1D | calcium/calmodulin-dependent protein kinase ID | 7.14E-06 |
| rs10906168 | 10 | 12597796 | 12597796 | intron variant; non-coding transcript variant | CAMK1D | calcium/calmodulin-dependent protein kinase ID | 7.14E-06 |
| rs10906170 | 10 | 12598308 | 12598308 | intron variant; non-coding transcript variant | CAMK1D | calcium/calmodulin-dependent protein kinase ID | 7.14E-06 |
| rs10906171 | 10 | 12602320 | 12602320 | intron variant; non-coding transcript variant | CAMK1D | calcium/calmodulin-dependent protein kinase ID | 7.14E-06 |
| rs10906173 | 10 | 12605105 | 12605105 | intron variant; non-coding transcript variant | CAMK1D | calcium/calmodulin-dependent protein kinase ID | 7.14E-06 |
| rs12764878 | 10 | 12603808 | 12603808 | intron variant; non-coding transcript variant | CAMK1D | calcium/calmodulin-dependent protein kinase ID | 7.14E-06 |
| rs2296754 | 10 | 12597108 | 12597108 | intron variant; non-coding transcript variant | CAMK1D | calcium/calmodulin-dependent protein kinase ID | 7.14E-06 |
| rs2648711 | 10 | 12605090 | 12605090 | intron variant; non-coding transcript variant | CAMK1D | calcium/calmodulin-dependent protein kinase ID | 7.14E-06 |
| rs77207912 | 15 | 45789759 | 45789759 | intron variant | SLC30A4 | solute carrier family 30 (zinc transporter), member 4 | 0.000135 |
| rs139638763 | 14 | 73821393 | 73821393 | intron variant; non-coding transcript variant | NUMB | numb homolog (Drosophila) | 0.000135 |
| rs34344194 | 12 | 15925995 | 15925995 | intron variant; NMD transcript variant | EPS8 | epidermal growth factor receptor pathway substrate 8 | 0.000135 |
| rs61908138 | 12 | 15946956 | 15946956 | intron variant | EPS8 | epidermal growth factor receptor pathway substrate 8 | 0.000135 |
| rs61908155 | 12 | 16010104 | 16010104 | intron variant | EPS8 | epidermal growth factor receptor pathway substrate 8 | 0.000135 |
| rs11159008 | 14 | 25431791 | 25431791 | intron variant; NMD transcript variant | STXBP6 | syntaxin binding protein 6 (amisyn) | 0.000283 |
| rs17257661 | 14 | 25432870 | 25432870 | intron variant; NMD transcript variant | STXBP6 | syntaxin binding protein 6 (amisyn) | 0.000283 |
| rs8181975 | 14 | 25433327 | 25433327 | intron variant; NMD transcript variant | STXBP6 | syntaxin binding protein 6 (amisyn) | 0.000283 |
| rs35691615 | 16 | 6495078 | 6495078 | intron variant; non-coding transcript variant | RBFOX1 | RNA binding protein, fox-1 homolog (C. elegans) 1 | 0.000305 |
| rs11041493 | 11 | 7664710 | 7664710 | intron variant; non-coding transcript variant | PPFIBP2 | PTPRF interacting protein, binding protein 2 (liprin beta 2) | 0.000552 |
| rs9424165 | 10 | 12607024 | 12607024 | intron variant; non-coding transcript variant | CAMK1D | calcium/calmodulin-dependent protein kinase ID | 0.000654 |
| rs378359 | 6 | 5123453 | 5123453 | intron variant; non-coding transcript variant; NMD transcript variant | LYRM4 | LYR motif containing 4 | 0.000718 |
| rs385481 | 6 | 5125574 | 5125574 | intron variant; non-coding transcript variant; NMD transcript variant | LYRM4 | LYR motif containing 4 | 0.000718 |
| rs10504272 | 8 | 59900124 | 59900124 | intron variant | TOX | thymocyte selection-associated high mobility group box | 0.001697 |
| rs111937104 | 8 | 59897223 | 59897223 | intron variant | TOX | thymocyte selection-associated high mobility group box | 0.001697 |

| | | | | | | | |
|---|---|---|---|---|---|---|---|
| rs61514480 | 8 | 59909111 | 59909111 | intron variant | TOX | thymocyte selection-associated high mobility group box | 0.001697 |
| rs78300448 | 8 | 59906910 | 59906910 | intron variant | TOX | thymocyte selection-associated high mobility group box | 0.001697 |
| rs2329670 | 5 | 41349485 | 41349485 | intron variant | PLCXD3 | phosphatidylinositol-specific phospholipase C, X domain containing 3 | 0.001774 |
| rs7774359 | 6 | 5486254 | 5486254 | intron variant | FARS2 | phenylalanyl-tRNA synthetase 2, mitochondrial | 0.002228 |
| rs75590771 | 2 | 171332206 | 171332206 | intron variant; non-coding transcript variant; NMD transcript variant | MYO3B | myosin IIIB | 0.002324 |
| rs141575794 | 17 | 62507740 | 62588982 | intron variant; NMD transcript variant | CEP95 | centrosomal protein 95kDa | 0.005114 |
| rs190729287 | 4 | 16050206 | 16050206 | intron variant; NMD transcript variant | PROM1 | prominin 1 | 0.005114 |
| rs138952073 | 5 | 26974554 | 26974554 | intron variant; non-coding transcript variant | CDH9 | cadherin 9, type 2 (T1-cadherin) | 0.005114 |
| rs146115003 | 14 | 93276818 | 93276818 | intron variant; non-coding transcript exon variant | GOLGA5 | golgin A5 | 0.005114 |
| rs12188035 | 5 | 76638444 | 76638444 | intron variant | PDE8B | phosphodiesterase 8B | 0.005114 |
| rs11621072 | 14 | 32908078 | 32908078 | intron variant; non-coding transcript exon variant | AKAP6 | A kinase (PRKA) anchor protein 6 | 0.005752 |
| rs1957014 | 14 | 32907086 | 32907086 | intron variant; non-coding transcript variant | AKAP6 | A kinase (PRKA) anchor protein 6 | 0.005752 |
| rs1006740 | 2 | 230471160 | 230471160 | intron variant | DNER | delta/notch-like EGF repeat containing | 0.007503 |
| rs10166515 | 2 | 230460953 | 230460953 | intron variant | DNER | delta/notch-like EGF repeat containing | 0.007503 |
| rs10176645 | 2 | 230463982 | 230463982 | intron variant | DNER | delta/notch-like EGF repeat containing | 0.007503 |
| rs10182101 | 2 | 230461604 | 230461604 | intron variant | DNER | delta/notch-like EGF repeat containing | 0.007503 |
| rs10933305 | 2 | 230468480 | 230468480 | intron variant | DNER | delta/notch-like EGF repeat containing | 0.007503 |
| rs11891200 | 2 | 230460267 | 230460267 | intron variant | DNER | delta/notch-like EGF repeat containing | 0.007503 |
| rs11891909 | 2 | 230471427 | 230471427 | intron variant | DNER | delta/notch-like EGF repeat containing | 0.007503 |
| rs12373570 | 2 | 230460640 | 230460640 | intron variant | DNER | delta/notch-like EGF repeat containing | 0.007503 |
| rs12373571 | 2 | 230460818 | 230460818 | intron variant | DNER | delta/notch-like EGF repeat containing | 0.007503 |
| rs12998709 | 2 | 230460466 | 230460466 | intron variant | DNER | delta/notch-like EGF repeat containing | 0.007503 |
| rs13021776 | 2 | 230460178 | 230460178 | intron variant | DNER | delta/notch-like EGF repeat containing | 0.007503 |
| rs1465346 | 2 | 230470702 | 230470702 | intron variant | DNER | delta/notch-like EGF repeat containing | 0.007503 |
| rs1465347 | 2 | 230470738 | 230470738 | intron variant | DNER | delta/notch-like EGF repeat containing | 0.007503 |
| rs1861616 | 2 | 230462275 | 230462275 | intron variant | DNER | delta/notch-like EGF repeat containing | 0.007503 |
| rs2052300 | 2 | 230461647 | 230461647 | intron variant | DNER | delta/notch-like EGF repeat containing | 0.007503 |
| rs2052303 | 2 | 230461850 | 230461850 | intron variant | DNER | delta/notch-like EGF repeat containing | 0.007503 |
| rs2052304 | 2 | 230462056 | 230462056 | intron variant | DNER | delta/notch-like EGF repeat containing | 0.007503 |
| rs2052306 | 2 | 230464359 | 230464359 | intron variant | DNER | delta/notch-like EGF repeat containing | 0.007503 |
| rs2075256 | 2 | 230462384 | 230462384 | intron variant | DNER | delta/notch-like EGF repeat containing | 0.007503 |
| rs2075257 | 2 | 230462394 | 230462394 | intron variant | DNER | delta/notch-like EGF repeat containing | 0.007503 |
| rs2396694 | 2 | 230462929 | 230462929 | intron variant | DNER | delta/notch-like EGF repeat containing | 0.007503 |

| | | | | | | | |
|---|---|---|---|---|---|---|---|
| rs2396695 | 2 | 230462932 | 230462932 | intron variant | DNER | delta/notch-like EGF repeat containing | 0.007503 |
| rs6751880 | 2 | 230463620 | 230463620 | intron variant | DNER | delta/notch-like EGF repeat containing | 0.007503 |
| rs759534 | 2 | 230462463 | 230462463 | intron variant | DNER | delta/notch-like EGF repeat containing | 0.007503 |
| rs759535 | 2 | 230463103 | 230463103 | intron variant | DNER | delta/notch-like EGF repeat containing | 0.007503 |
| rs759536 | 2 | 230463185 | 230463185 | intron variant | DNER | delta/notch-like EGF repeat containing | 0.007503 |
| rs759537 | 2 | 230463271 | 230463271 | intron variant | DNER | delta/notch-like EGF repeat containing | 0.007503 |
| rs759538 | 2 | 230463394 | 230463394 | intron variant | DNER | delta/notch-like EGF repeat containing | 0.007503 |
| rs759539 | 2 | 230465606 | 230465606 | intron variant | DNER | delta/notch-like EGF repeat containing | 0.007503 |
| rs888176 | 2 | 230461255 | 230461255 | intron variant | DNER | delta/notch-like EGF repeat containing | 0.007503 |
| rs888177 | 2 | 230461323 | 230461323 | intron variant | DNER | delta/notch-like EGF repeat containing | 0.007503 |
| rs1409961 | 10 | 12608676 | 12608676 | intron variant; non-coding transcript variant | CAMK1D | calcium/calmodulin-dependent protein kinase ID | 0.008404 |
| rs6602598 | 10 | 12606465 | 12606465 | intron variant; non-coding transcript variant | CAMK1D | calcium/calmodulin-dependent protein kinase ID | 0.008404 |
| rs7896236 | 10 | 12606281 | 12606281 | intron variant; non-coding transcript variant | CAMK1D | calcium/calmodulin-dependent protein kinase ID | 0.008404 |
| rs2037107 | 2 | 11299473 | 11299473 | intron variant; non-coding transcript variant; NMD transcript variant | PQLC3 | PQ loop repeat containing 3 | 0.008506 |
| rs11103520 | 9 | 137664056 | 137664056 | intron variant | COL5A1 | collagen, type V, alpha 1 | 0.013407 |
| rs6587924 | 1 | 61895257 | 61895257 | intron variant | NFIA | nuclear factor I/A | 0.015764 |
| rs6587925 | 1 | 61895589 | 61895589 | intron variant | NFIA | nuclear factor I/A | 0.015764 |
| rs4915741 | 1 | 61897148 | 61897148 | intron variant | NFIA | nuclear factor I/A | 0.019075 |
| rs62170724 | 2 | 171353456 | 171353456 | intron variant; non-coding transcript variant; NMD transcript variant | MYO3B | myosin IIIB | 0.023902 |
| rs4668271 | 2 | 171387405 | 171387405 | intron variant; non-coding transcript variant; NMD transcript variant | MYO3B | myosin IIIB | 0.024377 |
| rs62183592 | 2 | 190429752 | 190429752 | intron variant | SLC40A1 | solute carrier family 40 (iron-regulated transporter), member 1 | 0.026275 |
| rs2103883 | 1 | 61877378 | 61877378 | intron variant | NFIA | nuclear factor I/A | 0.029856 |
| rs9436640 | 1 | 61873677 | 61873677 | intron variant | NFIA | nuclear factor I/A | 0.029856 |
| rs1978315 | 16 | 6337331 | 6337331 | intron variant; non-coding transcript variant | RBFOX1 | RNA binding protein, fox-1 homolog (C. elegans) 1 | 0.032606 |
| rs117001634 | 10 | 24567837 | 24567837 | intron variant | KIAA1217 | KIAA1217 | 0.03341 |
| rs1108432 | 2 | 171368542 | 171368542 | intron variant; non-coding transcript variant; NMD transcript variant | MYO3B | myosin IIIB | 0.036459 |
| rs12881167 | 14 | 32908595 | 32908595 | intron variant | AKAP6 | A kinase (PRKA) anchor protein 6 | 0.037154 |
| rs2383304 | 14 | 32917431 | 32917431 | intron variant | AKAP6 | A kinase (PRKA) anchor protein 6 | 0.037154 |
| rs34976846 | 14 | 32916980 | 32916980 | intron variant | AKAP6 | A kinase (PRKA) anchor protein 6 | 0.037154 |
| rs186349113 | 11 | 18261841 | 18261841 | intron variant; non-coding transcript variant; splice_region_variant | SAA2-SAA4 | serum amyloid A2 | 0.038975 |
| rs113228935 | 3 | 155161226 | 155161226 | intron variant | PLCH1 | phospholipase C, eta 1 | 0.041618 |
| rs139867572 | 3 | 71534682 | 71534682 | intron variant; NMD transcript variant | FOXP1 | forkhead box P1 | 0.041618 |
| rs147398711 | 3 | 71526676 | 71526676 | intron variant; NMD transcript variant | FOXP1 | forkhead box P1 | 0.041618 |

| | | | | | | | |
|---|---|---|---|---|---|---|---|
| rs544840223 | 15 | 73505945 | 73505945 | intron variant | NEO1 | neogenin 1 | 0.041618 |
| rs17236331 | 5 | 68683069 | 68683069 | intron variant; non-coding transcript variant; NMD transcript variant | RAD17 | RAD17 homolog (S. pombe) | 0.041618 |
| rs10189774 | 2 | 171375497 | 171375497 | intron variant; non-coding transcript variant; NMD transcript variant | MYO3B | myosin IIIB | 0.044626 |
| rs10195667 | 2 | 171376832 | 171376832 | intron variant; non-coding transcript variant; NMD transcript variant | MYO3B | myosin IIIB | 0.044626 |
| rs7585843 | 2 | 171385679 | 171385679 | intron variant; non-coding transcript variant; NMD transcript variant | MYO3B | myosin IIIB | 0.044626 |
| rs7595366 | 2 | 171376648 | 171376648 | intron variant; non-coding transcript variant; NMD transcript variant | MYO3B | myosin IIIB | 0.044626 |
| rs1572687 | 13 | 55748673 | 55748673 | non-coding transcript exon variant | MIR5007 | microRNA 5007 | 0.046585 |

**Table S5. Transcript annotations for RNA transcripts determined in week 20 gestation analysis.** (bp=base pairs)

| Transcript cluster ID | Chromosome | Start position (bp) | End position (bp) | Gene | Gene name | Locus type | p |
|---|---|---|---|---|---|---|---|
| TC1000009465.hg.1 | chr10 | 133432816 | 133433528 | klernoby | Transcript Identified by AceView | Coding | 1.12E-06 |
| TC0100012239.hg.1 | chr1 | 244952545 | 244953016 | | | NonCoding | 6.48E-06 |
| TC0900009097.hg.1 | chr9 | 133849918 | 133850381 | gytarby | Transcript Identified by AceView | Coding | 3.01E-05 |
| TC1200009420.hg.1 | chr12 | 130953907 | 131141469 | ADGRD1 | adhesion G protein-coupled receptor D1 | Multiple_Complex | 4.95E-05 |
| TC1900007132.hg.1 | chr19 | 13298434 | 13298888 | nimoma | Transcript Identified by AceView | Coding | 5.59E-05 |
| TC0800012165.hg.1 | chr8 | 143790920 | 143815379 | SCRIB | scribbled planar cell polarity protein | Multiple_Complex | 5.72E-05 |
| TC0700008043.hg.1 | chr7 | 75267909 | 75268849 | parbley | Transcript Identified by AceView | Unassigned | 6.82E-05 |
| TC1400009036.hg.1 | chr14 | 42657763 | 42657918 | RP11-90P16.2 | | Multiple_Complex | 7.53E-05 |
| TC2000006635.hg.1 | chr20 | 6129526 | 6194558 | | | NonCoding | 7.98E-05 |
| TC1000009476.hg.1 | chr10 | 133576891 | 133577605 | lobawbu | Transcript Identified by AceView | Coding | 8.46E-05 |
| TC0700006476.hg.1 | chr7 | 926807 | 927368 | chychyby | Transcript Identified by AceView | Coding | 8.60E-05 |
| TC1700009262.hg.1 | chr17 | 82538654 | 82539436 | rorseebo | Transcript Identified by AceView | Coding | 8.85E-05 |
| TC2200008810.hg.1 | chr22 | 41065554 | 41065646 | Y_RNA | Y RNA | NonCoding | 9.45E-05 |

**Table S6. Metabolite bins (n=30) identified in week 20 gestation ANOVA with Tukey's HSD analysis (p<0.05).**

Breakdown of metabolite bins with p<0.05 for each phenotypic group comparison: PPROM-HTERM (n=0); SPTB-HTERM (n=28); SPTB-PPROM (n=5).

| Omic variables | ANOVA | PPROM-HTERM | SPTB-HTERM | SPTB-PPROM |
|---|---|---|---|---|
| 2-hydroxyvalerate/Arginine (1.62ppm) | 0.005981985 | 0.317407082 | 0.004745062 | 0.396507529 |
| acetoacetate (2.23ppm) | 0.006504984 | 0.181966441 | 0.006613724 | 0.628411825 |
| unknown (3.67ppm) | 0.011504893 | 0.484905755 | 0.039531448 | 0.014043317 |
| unknown (3.64ppm) | 0.011797459 | 0.495536833 | 0.039436656 | 0.014582383 |
| unknown (3.64ppm) | 0.011923646 | 0.491242963 | 0.040266434 | 0.014605515 |
| unknown (3.81ppm) | 0.012225832 | 0.988025277 | 0.013123946 | 0.050913588 |
| mannose (5.19ppm) | 0.012395051 | 0.189637433 | 0.013872525 | 0.749543263 |
| glycylproline (2.05ppm) | 0.013377343 | 0.685718427 | 0.009633989 | 0.231211113 |
| proline (2.33ppm) | 0.01385456 | 0.665880446 | 0.010004117 | 0.246838666 |
| creatinine (4.06ppm) | 0.017314103 | 0.33298294 | 0.015424592 | 0.583608344 |
| Lactate (1.33ppm) | 0.022553623 | 0.23666282 | 0.025029697 | 0.79002888 |
| 2-hydroxyvalerate (4.07ppm) | 0.022824868 | 0.498021122 | 0.018090721 | 0.454670731 |
| unknown (4.32ppm) | 0.024448161 | 0.986712106 | 0.026243367 | 0.079976945 |
| mobile-lipids (0.90ppm) | 0.026499959 | 0.483492918 | 0.02155069 | 0.499312719 |
| unknown (3.07ppm) | 0.026795114 | 0.148783525 | 0.304060848 | 0.020099194 |
| unknown (3.88ppm) | 0.02720249 | 0.989458759 | 0.028798359 | 0.087826648 |
| unknown (3.00ppm) | 0.029187566 | 0.554918074 | 0.081300188 | 0.033281209 |
| glutamate (2.48ppm) | 0.029588362 | 0.455152886 | 0.024912439 | 0.552852743 |
| glutamate (2.26ppm) | 0.030311325 | 0.549319901 | 0.024041738 | 0.463276607 |
| glucose (3.90ppm) | 0.031075053 | 0.908908071 | 0.024637055 | 0.202611324 |
| glucose (3.86ppm) | 0.031388565 | 0.866276128 | 0.024321819 | 0.231857047 |
| myoinositol (3.58ppm) | 0.032249443 | 0.999700981 | 0.03147585 | 0.114291223 |
| unknown (2.44ppm) | 0.036772051 | 0.997864952 | 0.036818664 | 0.118243598 |
| mobile-lipids (1.29ppm) | 0.038035222 | 0.40938044 | 0.034659561 | 0.663003579 |
| unknown (1.37ppm) | 0.03810343 | 0.313455786 | 0.039860132 | 0.788220327 |
| glutamate/proline (2.09ppm) | 0.038190517 | 0.989709231 | 0.034002485 | 0.157076239 |
| unknown (3.80ppm) | 0.038251347 | 0.80728042 | 0.029433878 | 0.300351762 |
| glucose (3.79ppm) | 0.041571769 | 0.816916078 | 0.032182452 | 0.306615448 |
| unknown (1.41ppm) | 0.044189772 | 0.858027129 | 0.034649166 | 0.28583069 |
| unknown (1.54ppm) | 0.048871908 | 0.464094609 | 0.043666453 | 0.658250889 |

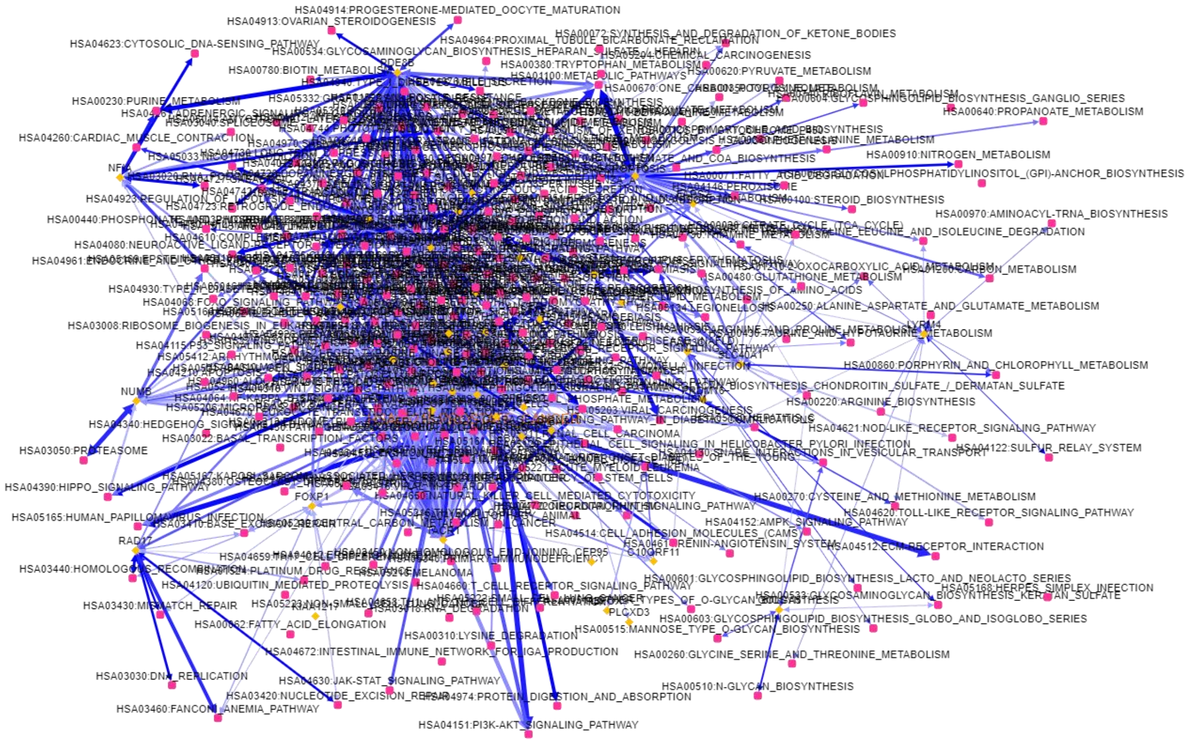


**Figure S1. Week 16 network of significant ANOVA genes generated using EviNet** (Jeggari et al., 2018). Yellow dot = input gene set; pink dots = functional gene sets; orange dots = both input and functional gene set; Two-headed arrows: gene-gene links, where edge transparency is confidence of network enrichment and edge thickness stands for number of gene-gene links.

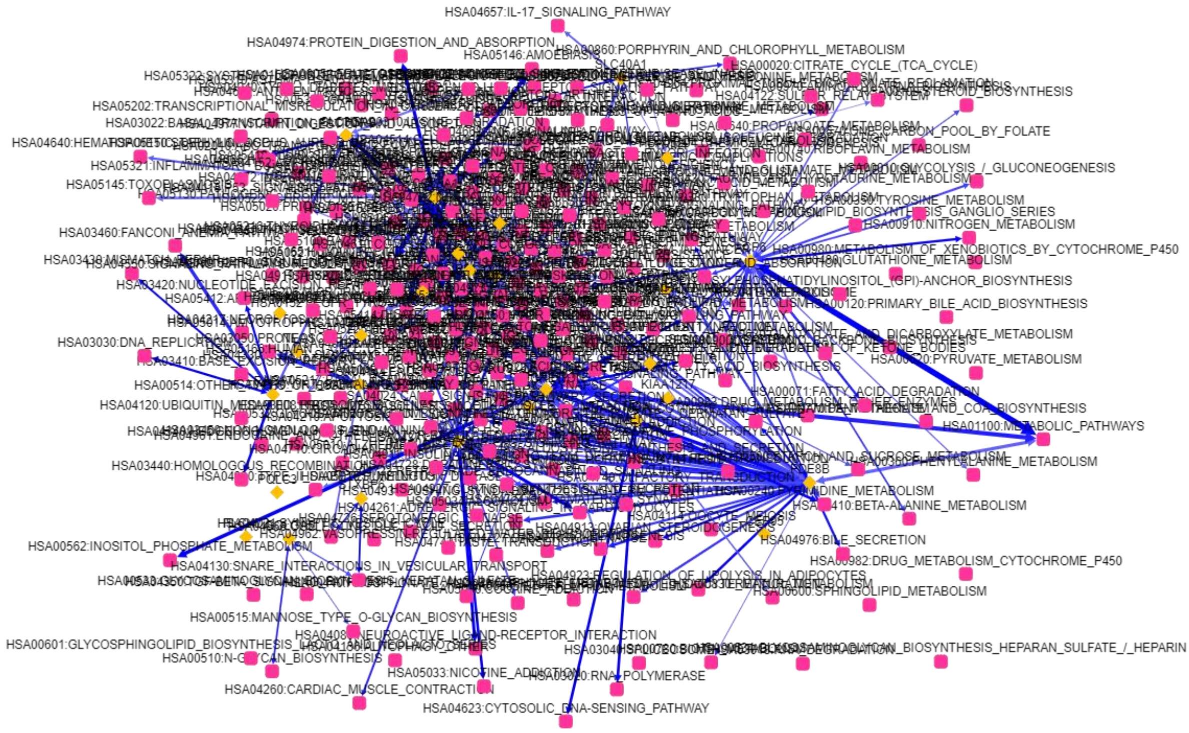


**Figure S2. Week 20 network of significant ANOVA genes generated using EviNet** (Jeggari et al., 2018). Yellow dot = input gene set; pink dot = functional gene sets; orange dots = both input and functional gene set; Two-headed arrows: gene-gene links, where edge transparency is confidence of network enrichment and edge thickness stands for number of gene-gene links.
